# Restoration of ovarian endocrine function with encapsulated immune isolated human ovarian xenograft in ovariectomized mice

**DOI:** 10.1101/2025.02.27.640622

**Authors:** Margaret A. Brunette, Monica A. Wall, Delaney Sinko, Jordan H. Machlin, Gabrielle M. Blevins, Marisa Leo, Ansen Tan, Bipasha Ray, Marilia Cascalho, Vasantha Padmanabhan, Ariella Shikanov

## Abstract

Anti-cancer treatments cause premature depletion of the non-renewable ovarian reserve of follicles, the source of key steroid hormones, leading to premature ovarian insufficiency (POI) in 50% of pediatric cancer survivors. Patients with POI, especially at the onset of pubertal development, experience significant endocrine complications, including delayed growth, elevated risks of obesity and diabetes, and accelerated cardiovascular, musculoskeletal and neurological disorders as adults. The only approved pharmacological treatment for POI is an off-label prescribed hormone replacement therapy, which does not replace physiologically functioning ovaries. To restore production of ovarian hormones and protect against immune-mediated injury, we developed a hydrogel-based capsule for implantation of donor ovarian tissue. We evaluated the restoration of ovarian endocrine function in ovariectomized immunodeficient (NOD scid gamma, NSG) mice implanted with encapsulated xenografts over 20 weeks through daily vaginal cytology, hormone measurements and histological analysis of explanted human xenografts. The encapsulated xenografts integrated into the murine hypothalamus-pituitary-gonad (HPG) axis responding to circulating murine gonadotropins and restoring ovarian endocrine function. As controls, we implanted non encapsulated human ovarian xenografts comparable in size. Without the need for exogeneous stimulation, the estrous cyclicity resumed in both groups of mice 12 weeks post implantation and all mice regularly cycled experiencing between 3 to 8 estrous cycles in 20 weeks. The levels of estradiol gradually increased reaching on average 50pg/mL 20 weeks post implantation. Morphological analysis of the encapsulated grafts revealed presence of large antral follicles, ∼3mm in diameter, consistent with regular cyclicity and measurable levels of circulating hormones. This work demonstrates that endocrine function of encapsulated human ovarian tissue was not affected by the encapsulation and integrated with the host physiology similarly to the non-encapsulated controls.

## 1. Introduction

The population of pediatric cancer survivors continues to grow with an estimated 9,000 new diagnoses in 2024 for children aged 0-14(*1, 2*). This results in more children and teenagers being diagnosed with and surviving cancer, and living into adolescence and adulthood(*1, 2*). While having cancer in remission is a significant milestone, it is important to consider the patient’s quality of life after treatment. Chemotherapy and radiation are particularly harmful to the nonrenewable follicular reserve in the ovaries(*3*). Damage to the ovaries may prevent or delay development in these survivors, with 50% of this population experiencing health complications and infertility(*3–6*). The lack of ovarian hormones leads to a cascade of other negative outcomes including cardiac disease, osteopenia, growth hormone deficiency, loss of neuroprotection, as well as dysregulated T cell function, premature epiphyseal closure, and imbalance in food intake and serum leptin levels(*7–12*). The widespread systemic impact of the lost ovarian endocrine function demonstrates the importance of ovaries and its hormonal feedback in female pediatric cancer survivors.

Hormone replacement therapy (HRT) is the only available treatment to induce puberty in adolescent patients in remission who experience premature ovarian insufficiency (POI)(*13, 14*). Using HRT for puberty induction in adolescent girls with POI has multiple limitations(*15–19*). First and most importantly, HRT supplies only two hormones, estradiol and progesterone, at fixed levels lacking the pulsatile and cyclic fluctuations normally present during the circadian and menstrual cycles(*16–18*). Second, the levels of hormones delivered with HRT fail to mimic the reciprocal physiological feedback systems in the hypothalamic-pituitary gonadal (HPG) axis and the rest of the body. Third, the exogenous non-physiological pattern of delivery of steroid hormones has been associated with higher risk of anxiety and suicide, particularly for adolescents(*19*). Lastly, the long-term safety and efficacy of HRT for puberty induction has yet to be evaluated in adolescent girls requiring the HRT treatment for decades to compensate for the loss of gonadal steroids.^18^

As an alternative for HRT, implanted donor ovarian tissue offers a viable approach to restore the lost ovarian endocrine milieu and the cyclical reciprocal feedback of ovarian hormones to the HPG axis. In the past decade implantation of autologous human ovarian cortex has been successful for fertility restoration and can be offered to pediatric patients and women unable to undergo ovarian stimulation(*20, 21*). Fresh ovarian tissue transplantation from an identical twin was first shown to be successful in 2005 with documented restoration of fertility and endogenous endocrine function(*22*). Transplantation of cryopreserved autologous tissue restored estrogen production in 75% of participants, decreased follicle stimulating hormone (FSH) in 72% of participants, decreased luteinizing hormone (LH) in 67% of participants, and led to the return of menses in 72% of participants(*22*). Pooling data from all participants (n=568 women), regardless of the graft being fresh or cryopreserved, Khattak *et al*. concluded that FSH and LH returned to physiological levels 19 weeks after implantation, and menses returning after 18 weeks(*22*). However, the use of autologous tissues for pediatric survivors carries the risk of reintroducing cancer cells potentially harbored in the tissue, especially in patients with hematologic cancers. Furthermore, this approach is limited to patients who cryopreserved their tissue prior to gonadotoxic treatment, which is far from a widespread practice. In contrast, immune-isolated donor ovarian tissue offers a safe and accessible approach to restore physiological cyclic ovarian endocrine function without immune suppression.

Human ovarian tissue has been successfully xenografted and shown to survive from days to months in ovariectomized or intact immunocompromised hosts. Many different approaches were tested, including stimulation of folliculogenesis in the implanted tissue with exogeneous FSH, varying the location of the grafts (intraperitoneal, subcutaneous, intramuscular, and in ovarian bursa), use of immunocompromised mice such as the NSG mice that have the *Prkdc* variant associated with severe combined immunodeficiency and a knockout of the IL-2 receptor gamma chain gene resulting in a deficiencies in B and T lymphocytes and NK cells; and congenitally athymic nude mice (nu/nu)(*23*). Yet, implantation of a non-genetically identical donor tissue would require immune suppression to prevent rejection. We hypothesized that encapsulating donor ovarian tissue in a hydrogel-based immune isolating capsule would allow bidirectional diffusion of nutrients and hormones while preventing immune rejection. Towards that hypothesis, we have developed an immune-isolating capsule that supported implantation of allogeneic murine tissue in immunocompetent hosts for weeks, after repeated rounds of implantations in naïve and immune sensitized rodent hosts. Our previous work demonstrated that a poly(ethylene glycol) (PEG)-based hydrogel capsule with a degradable core and a non-degradable shell supported growth and function of murine follicles in the encapsulated allogeneic tissue while precluding immune cell infiltration and rejection of the allograft(*24, 25*). We also demonstrated that encapsulated 1mm^3^ human ovarian tissues survived in immunocompromised mice for a month (*26*). However, to extend graft longevity and implant a greater number of follicles heterogeneously distributed within the human ovarian cortex we designed larger immune-isolating capsules. In this investigation (**Figure 1A**), we increased the volume of the graft from 1mm^3^ to 19.6mm^3^ and tested whether encapsulated human follicles resumed folliculogenesis, reached antral stages, integrated with mouse HPG axis, and secreted gonadal hormones in ovariectomized mice. We found over a 20-week period that (1) estrous cyclicity resumed in all mice (2) estradiol levels increased significantly with graft implantation, and (3) FSH levels trended downward after 18 weeks. Morphological analysis of the encapsulated grafts confirmed the presence of large antral follicles, ∼3mm in diameter, responsible for the hormone production and similar to non-encapsulated controls. This work demonstrates, for the first time that the survival and function, including hormone production and folliculogenesis, of encapsulated human ovarian tissue was not affected by the lack of direct vascularization. Furthermore, we demonstrated that encapsulated human ovarian tissue integrated into the mouse HPG axis, responding to circulating host’s gonadotropins and secreting estradiol similar to non-encapsulated controls.

**Figure 1:**
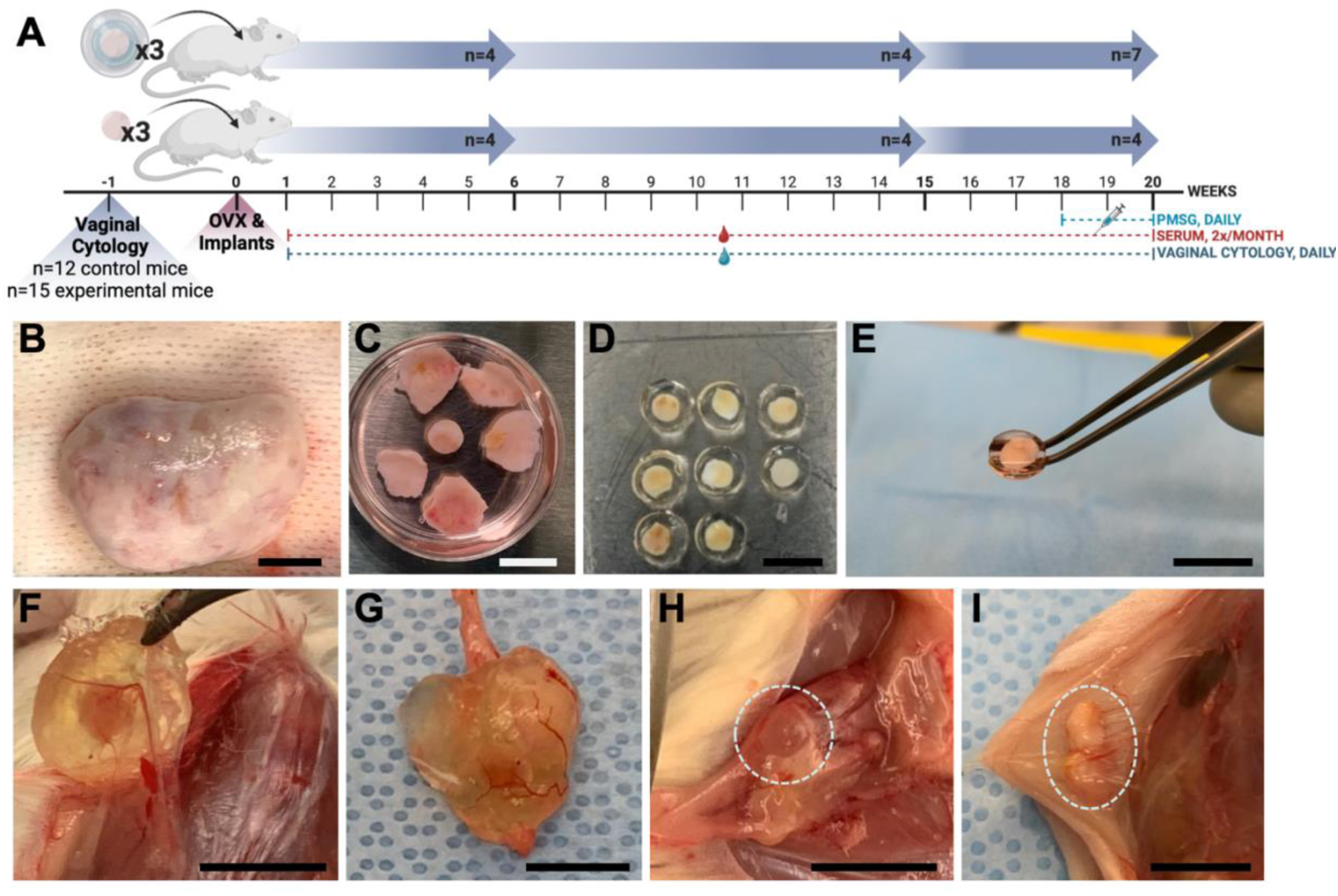
NSG mice were implanted with human cortical ovarian tissue (encapsulated or non­encapsulated) and followed out to twenty weeks (A). Donor ovaries (B) are de-cortified (C) and cryopreserved until use. After thawing, tissue is encapsulated in the immune-isolating capsule (D) and is implanted (E). Encapsulated tissue samples (F,G) and non-encapsulated tissue samples (HJ) were removed from the subcutaneous space. Panels G and H show gross morphological changes indicating antral follicles are present. Scale bar represents 1cm A-G, and 5mm for H & I.

## 2. Materials and Methods

### 2.1 Collection of Human Ovarian Tissue

Organ procurement was conducted as previously described(*26*). Briefly, ovaries from healthy reproductive age women were obtained following standardized protocols of the International Institute for the Advancement of Medicine (IIAM) and the associated Organ Procurement Organization (OPO) for research purposes. This study utilized ovaries procured by the IIAM from eight deceased donors ranging between 16-31 years of age. Organs were placed in storage solution and shipped on ice. Storage solutions were either normal saline or Belzer University of Wisconsin® Cold Storage Solution. **Supplemental Table 1** summarizes de-identified donor information namely age, height, weight, BMI, right and left ovarian volume, cardiac arrest/downtime, cross-clamp time (time at which the organ is cut from blood/oxygen supply), cold ischemic time, ethnicity, and cause of death. Cold ischemic time (CIT) indicates the time interval between cross-clamp time of the donor (and subsequent cessation of arterial blood flow to the ovaries) in the operation room and start time of the tissue harvest after arrival at the laboratory.

### 2.2 Ethical Approval Process

As described previously(*26*), the IIAM procures tissue and organs for non-clinical research from Organ Procurement Organizations (OPOs), which comply with state Uniform Anatomical Gift Acts (UAGA) and are certified and regulated by the Centers for Medicare and Medicaid Services (CMS). These OPOs are members of the Organ Procurement and Transplantation Network (OPTN) and the United Network for Organ Sharing (UNOS) and operate under a set of standards established by the Association of Organ Procurement Organizations (AOPO) and UNOS. Informed, written consent from the deceased donor’s family was obtained for the tissue used in this publication. A biomaterial transfer agreement is in place between IIAM and the authors that restricts the use of the tissue for pre-clinical research that does not involve the fertilization of gametes. The use of deceased donor ovarian tissue in this research is categorized as ‘not regulated’, per 45 CFR 46.102 and the ‘Common Rule’, as it does not involve human subjects and complies with the University of Michigan’s IRB requirements as such.

### 2.3 Tissue Processing

All tissue processing was done aseptically in a biosafety cabinet. After receiving donor tissues, the ovaries were first separated from other reproductive tissues (i.e. the uterus, fallopian tubes) (**Figure 1B**). The ovaries were de-cortified using a custom cutting guide (Reprolife Japan, Tokyo) to make squares measuring 10mm x 10mm and approximately 1 mm thick (**Figure 1C**). The squares were aseptically transferred into cryovials with 1mL holding media (Quinn’s Advantage Medium with HEPES (QAMH), 10% Quinn’s Advantage Serum Protein Substitute (SPS), CooperSurgical, Måløv, Denmark).

### 2.4 Slow Freezing

The methods described by Xu et al. were used for slow freezing(*27*). Briefly, the 10mm x 10mm x 1mm squares of cortical tissue were placed into cryovials (Nunc, Roskilde, Denmark) filled with pre-cooled cryoprotectant media (QAMH, 10% SPS, 0.75M dimethyl sulfoxide (DMSO) (Sigma Aldrich, St. Louis, USA), 0.75M ethylene glycol (Sigma Aldrich, St. Louis, USA), 0.1M sucrose (Sigma Aldrich, St. Louis, USA)), and equilibrated at 4°C for at least 30 minutes. After equilibration, cryovials were loaded into the Cryologic Freeze Control System (Cryologic, Victoria, Australia). Vials were then frozen via the following protocol: (1) cooling from 4°C to −9°C at a rate of −2°C /min (2) equilibration for 6 min at −9°C (3) manual seeding using large swabs cooled by submersion in liquid nitrogen (4) holding for 4 min at −9°C (5) cooling to −40°C at a rate of −0.3°C/min and (6) storage in liquid nitrogen a cryogenic storage dewar until thawed for use.

### 2.5 Thawing Tissue

The process described by Brunette et al. was followed with minimal changes(*26*). Briefly, vials with ovarian tissues were removed from liquid nitrogen and placed in a 37°C bath. Once the cryoprotectant media in the vial had thawed, the tissue was removed from the vial and put into Thaw Solution One (0.5M DMSO, 0.5M ethylene glycol, 0.1M Sucrose, 10% SPS in QAMH) for ten minutes. Tissue was then incubated sequentially in Thaw Solution Two (0.25M DMSO, 0.25M ethylene glycol, 0.1M Sucrose, 10% SPS in QAMH), Three (0.1M Sucrose, 10% SPS in QAMH), and Four (10% SPS in QAMH) for ten minutes each. Thaw solutions were maintained at room temperature, and samples were protected from light and agitated while in thaw solutions. Tissue squares were biopsy punched into disks measuring approximately 5mm in diameter while still in Thaw Solution Four (**Figure 1C**).

### 2.6 Encapsulation

The disks of ovarian cortical tissue (5mm diameter) were maintained in the final thaw solution media (Solution 4) at 37°C until encapsulation. The PEG core was prepared by cross-linking 8-arm PEG-VS (40 kDa, Jenkem Technology, Beijing, China) (5% w/v) with plasmin sensitive peptide (Ac-GCYK↓NSGCYK↓NSCG, MW 1525.69 g/mol, > 90% Purity, Genscript, ↓ indicates the cleavage site of the peptide). The PEG shell was prepared with 4-arm PEG-VS (20 kDa, Jenkem Technology) (5% w/v), Irgacure 2959 (Ciba, Switzerland, MW = 224.3) (0.4% w/v), and N-vinyl-2-pyrrolidone (Sigma-Aldrich, St. Louis, USA) (0.1% v/v).

The tissue was placed in a mold and 40µL of degradable PEG core pre-cursor solution was added, covering the tissue surface. After five to six minutes of crosslinking the core was removed from the mold and set on top of 70µL of non-degradable PEG shell pre-cursor solution. An additional 30-35µL of non-degradable PEG shell precursor solution was added on top of the core, ensuring complete coverage. The shell was cross-linked via UV light (OmniCure S1500, Excelitas Technologies, USA) at constant intensity (30mW/cm^2^) for 45 seconds. Encapsulated tissue (**Figure 1E**) was maintained in Leibovitz L-15 media (Gibco, USA) at 37°C until implantation.

### 2.7 Subcutaneous Implantation

Animal experiments for this work were performed in accordance with the protocol approved by the Institutional Animal Care and Use Committee (IACUC) at the University of Michigan (PRO00009635). The IACUC guidelines for survival surgery in rodents and the IACUC Policy on Analgesic Use in Animals Undergoing Surgery were followed for all procedures.

Female NSG mice (strain 005557, The Jackson Laboratory, Bar Harbor, ME, USA) 8-18 weeks old were anesthetized using isoflurane (2-3%) via inhalation. Mice were given preemptive analgesics (Carprofen, RIMADYL, Zoetis, USA, 5mg/kg body) via subcutaneous injection. An incision was made in the medial/dorsal skin. Bilateral ovariectomies were performed before graft implantation. Three PEG capsules with human cortical issue or three control non-encapsulated tissues were inserted into the dorsal subcutaneous space in the mouse. The control graft was inserted subcutaneously and sutured using 5/0 absorbable sutures (AD surgical) to the subcutaneous tissue to ensure graft recovery. Using 5/0 absorbable sutures the incision was closed, with special attention paid to avoid suturing capsules/control tissue. Mice recovered in a clean cage, were given analgesics at 24 hours, and were monitored post-operatively for 7-10 days.

### 2.8 Serum Hormone Analysis

Every two weeks blood was collected from the lateral tail vein. A volume no greater than 1% body weight was collected each time using glass Pasteur pipettes (FisherBrand). At the time of sacrifice, blood was collected via cardiac puncture. Following blood collection, samples were stored at 4 °C overnight, then centrifuged for 15 minutes at 2,000 G. Serum was aliquoted and stored at −80 °C until analysis. The samples were analyzed for mouse FSH and estradiol at the Ligand Assay and Analysis Core Facility at the University of Virginia Center for Research in Reproduction. FSH was analyzed using a radioimmunoassay, with a lower detection limit of 1.9 ng/mL. Estradiol was analyzed using ALPCO ELISA, with a lower detection limit of 5 pg/mL.

### 2.9 Vaginal Cytology

Vaginal cytology was collected daily starting 7 days post-surgery until completion of the study(*25, 28*). Briefly, sterile saline was used to collect a sample of cells present in the vaginal lavage and analyzed to determine the composition of cell types present. Each stage of the estrous cycle corresponds with a typical population of cells in the vaginal lavage: diestrus is characterized by a presence of small, round leukocytes; proestrus presents with nucleated epithelial cells; estrus is dominated by cornified epithelial cells followed with metestrus that presents with a mixed population of nucleated and cornified epithelial cells. The loss of estrous cyclicity after mice undergo ovariectomies was confirmed by a persistent diestrus with only leukocytes present in the vaginal lavage. The cyclical transition of cells from leukocytes to cornified epithelial and then to nucleated epithelial cells signaled resumption of estrous cyclicity.

### 2.10 PMSG Injections

Mice were stimulated with Pregnant Mare Serum Gonadotropins (PMSG, Pregnyl, Merck, USA) injections for 2 weeks starting on week 18 of the study. Mice were injected with 2IU PMSG each day via intraperitoneal injections until euthanasia at the end of the experiment at 20 weeks.

### 2.11 Implant Removal

Implants were removed at 6, 15, or 20 weeks. Mice were anesthetized using isoflurane as described above. An incision was made in the medial/dorsal skin, avoiding implanted grafts. The encapsulated (**Figure 1F, G**) and control grafts (**Figure 1H, I**) were removed and placed in 4% PFA overnight at 4°C, washed, and stored in 70% ethanol.

### 2.12 Histology

All samples were processed at the Histology Core in the Dental School at the University of Michigan. The paraffin embedded tissue blocks were serially sectioned at a thickness of 5µm and stained with hematoxylin and eosin.

### 2.13 Follicle Counting

Slides with ovarian tissue sections were imaged using a digital slide scanner (Leica Aperio AT2®, Germany) at 20x magnification. Every 8th section was analyzed for the number and stage of follicles present, with 15 to 25 sections analyzed per donor. Each data point represents the follicle density calculated from each section analyzed. QuPath software (v0.2.2) was used for manual follicle counting. Follicle stage was identified using standard morphological guidelines(*29*). All primordial and primary follicles were counted for each analyzed section. Preantral and antral follicles were counted only after comparing the location of the follicle in the tissue and follicle size in preceding and subsequent sections to avoid “double counting”. Density measurements were calculated using tissue section area and the estimated tissue section thickness (20µm). The follicle density values reported in this study are all normalized to the same volume.

### 2.14 Statistical Analysis

Statistical analysis was performed using GraphPad Prism software, version 10. To determine whether there was a significant difference between the number of estrus cycles between encapsulated and non-encapsulated groups a two-way ANOVA with Šídák’s multiple comparisons was used. Estradiol levels between ovariectomized controls, non-encapsulated, and encapsulated groups were compared with a one-way ANOVA with Tukey’s multiple comparisons tests. Estradiol levels between non-encapsulated and encapsulated groups at each time point were evaluated with a one-way ANOVA with multiple comparisons. For all test, results were considered statistically significant when p < 0.05.

## 3. Results

### 3.1 Morphometry of the donor ovarian tissue demonstrates inter- and intra-donor heterogeneity of follicle distribution

Representative histological images for each of the 8 donors used in this study documenting the presence of primordial follicles are shown in **Figure 2A-H**. Tissue from Donor A (18 years old (y.o.)), B (16 y.o.), and C (25 y.o.) qualitatively show a higher density of primordial follicles, which is supported by follicle density measurements (**Figure 2I**) with 662 ± 330, 297 ± 68, and 107 ± 98 follicles per mm^3^ for Donors A, B, and C, respectively. The follicle density for Donors D (27 y.o.), E (18 y.o.), F (25 y.o.), G (31 y.o.), and H (28 y.o.) was lower: 56 ± 35, 54 ± 54, 2 ± 3, 24 ± 15, and 50 ± 51 follicles per mm^3^ respectively. To compensate for differences in age and follicular density between available donors, tissue from a mix of donors (summarized in **Figure 2J, K**) were implanted into the same mouse recipient. The average age of donor tissue used was similar for all mice in all conditions: 24.0 ± 6.0 years (yrs.) for the non-encapsulated control mice (NE) and 23.0 ± 6.0 yrs. for mice receiving encapsulated ovarian tissue (E) for the 6-week cohort, 23.8 ± 5.8 yrs. (NE) and 23.6 ± 6.4 yrs. (E) for the 15-week cohort, and 22.2 ± 5.8 yrs. (NE) and 23.0 ± 6.2 yrs. (E) for the 20-week cohort. Additional details on which donors were implanted in individual mice can be found in **Supplemental Table 2**.

**Figure 2:**
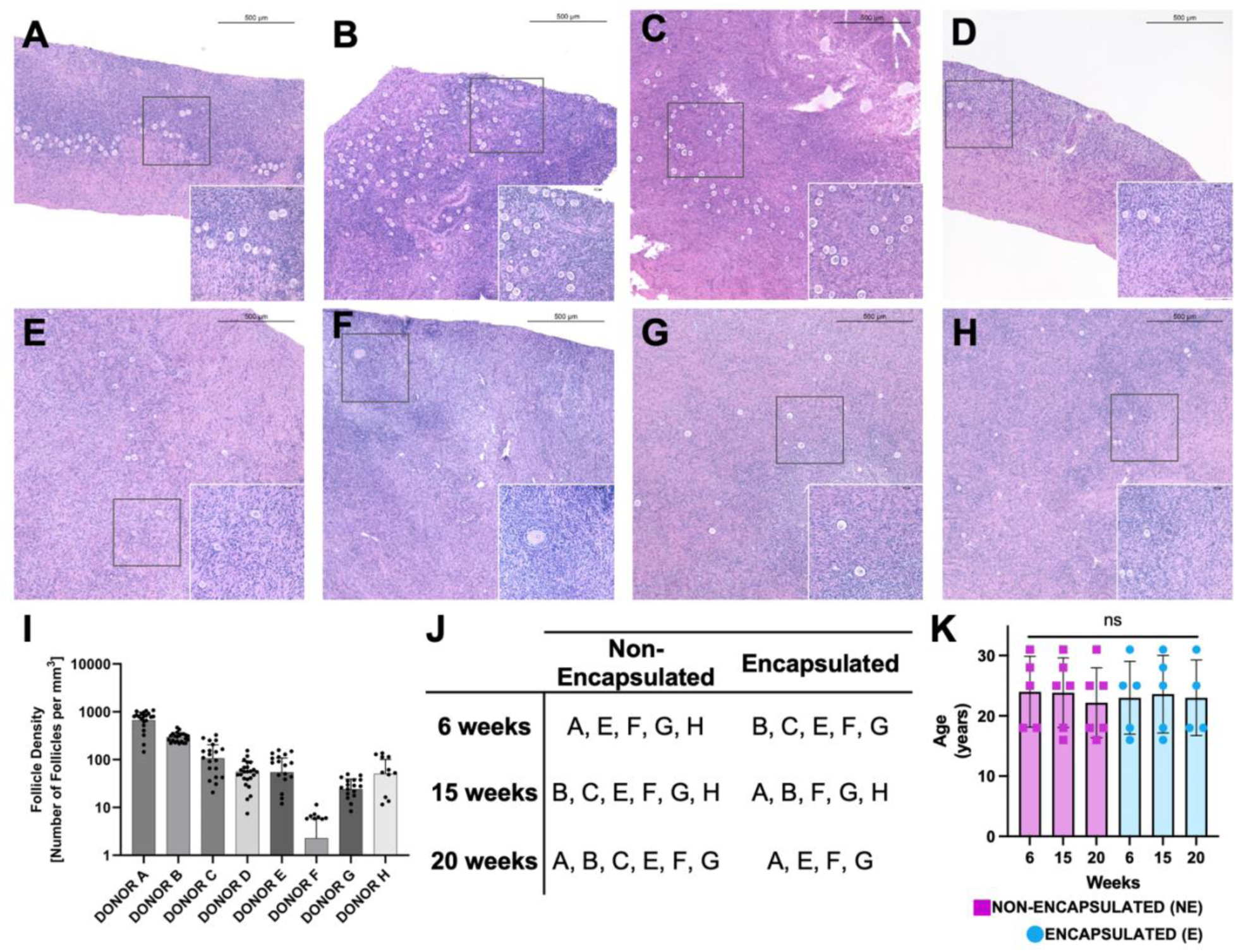
Representative histological images from fresh fixed human ovarian cortex from each donor utilized in this study is shown in A-H. The follicle densities for each donor are shown in I. The distribution of donor tissue among each group is shown in J, with the average age for each group shown in K.

### 3.2 Mice implanted with ovarian xenografts resumed estrous cyclicity

To determine graft functionality, we monitored vaginal opening and vaginal cytology daily. Following ovariectomy, all mice presented with closed vaginal openings, as expected, due to the lack of ovarian hormones. Changes in vaginal opening size appeared starting 4 weeks post implantation in mice with encapsulated and non-encapsulated human ovarian xenografts (**Figure 3A**). Next, to establish whether mice resumed estrous cyclicity we analyzed daily vaginal cytology for the proportion of cells types present (*30*) with a predominance of leukocytes signifying diestrus, or a prominence of cornified epithelial cells signifying estrus. Ovariectomized mice without ovarian function presented with persistent diestrus, while periodic changes in vaginal cells in mice who received implants suggested the production of required ovarian hormones by implanted grafts. Representative estrous cyclicity plots are shown in **Figure 3B** for a non-encapsulated human ovarian xenograft control and **Figure 3C** for mice implanted with encapsulated human tissue. Mice in both groups presented with multiple estrous peaks, followed with metestrus and diestrus, suggestive of resumption of regular cyclic ovarian function. The cycle frequencies and lengths varied between individual mice, possibly due to the heterogeneity of the human tissue. The first evidence of cyclicity was recorded 4 weeks post implantation with 20% of mice cycling in the control group and 38% in the encapsulated group (**Figure 3D**). By week six post implantation 70% in the control group and 62% in the test group cycled. By week eight, 90% in the control and 85% in the test group cycled. By week ten 90% in the control and 92% in the test group experienced at least one estrous cycle. By week twelve 100% of mice achieved estrus and continued cycling until they were euthanized at 20 weeks. (Detailed individual cyclicity data for each mouse are shown in Supplemental Figure 1). The number of cycles per group was analyzed for cohorts that were euthanatized at 15 and 20 weeks of the study. There was no statistical difference between the control non-encapsulated and test encapsulated groups in analyzed cohorts (**Figure 3E**), suggesting that encapsulation in the immune-isolating hydrogel does not impair hormone production of donor ovarian tissue. The average number of cycles for the 15-week cohort was 3.0 ± 1.8 for the control and 3.3 ± 1.5 for the test groups and the average number of cycles for the 20-week cohort was 3.5 ± 0.6 for the control and 4.9 ± 2.1 for the test groups. There were no significant differences in number of cycles between non-encapsulated and encapsulated groups.

**Figure 3:**
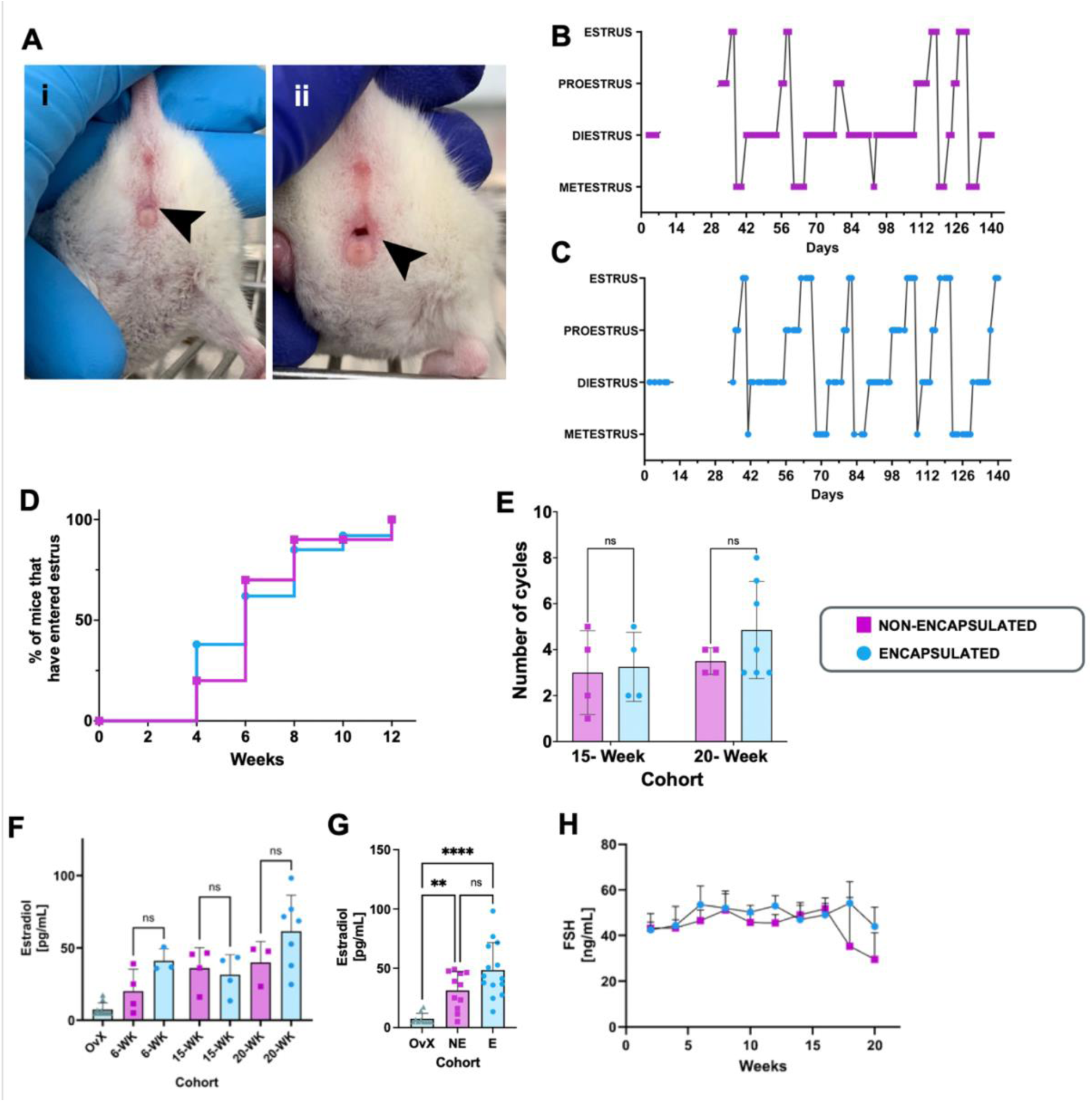
Endocrine function of the grafts is expected to be seen in four metrics: vaginal opening, estrous cyclicity, estradiol levels, and follicle stimulating hormone (FSH) levels, levels. Vaginal openings (black arrows) were found to be closed after ovariectomy (Ai), followed by opening (Aii). Vaginal opening was correlated with the resumption of estrus cyclicity. There is a resumption of cyclicity around five weeks for mice with non-encapsulated (B) and encapsulated (C) samples. Trends between non-encapsulated and encapsulated groups were similar when looking at the percent of mice that had entered estrus (D) and the number of cycles achieved in each cohort (F). Estradiol measurements were determined from terminal blood serum (F,G). Mice that received grafts showed statistically significant more estradiol than controls (F). Further analysis shows that there is no significant difference in estradiol levels between groups when comparing cohorts (G). Blood serum collected every other week shows a trend toward decreasing FSH levels after week 18 (H).

### 3.3 Human ovarian xenografts responded to endogenous murine gonadotropin stimulation by producing and secreting ovarian hormones as was confirmed by measurable levels of circulating estradiol

We hypothesized that by ovariectomizing graft recipients the lack of negative feedback from the native murine ovary would impact the levels of circulating FSH such that they would be high enough and sufficient to stimulate folliculogenesis and steroidogenesis in the human tissue without additional exogeneous stimulation with gonadotropins. After 15 weeks post implantation the levels of circulating estradiol reached 31.4 ± 13.9 pg/mL in mice implanted with encapsulated tissue and 36.1 ± 14.1 pg/mL in the control group implanted with non-encapsulated tissue (**Figure 3F**). After 20 weeks the estradiol levels reached 40.0 ± 14.5 pg/mL in the control (NE) and 61.5 ± 24.92 pg/mL in the test group (E), demonstrating continuous production and secretion of estradiol. These findings validate that endogenous murine FSH was able to diffuse through the capsule and stimulate estradiol production the encapsulated tissue to the same extent as the non-encapsulated tissue. The average levels of estradiol throughout the study were 48.5 ± 23.1 pg/mL for the test (E) group and 31.3 ± 15.9 pg/mL for the control group (NE), both were significantly higher than levels in ovariectomized or intact control mice (7.5 ± 4.5 pg/mL). There was no significant difference between the levels of estradiol measure in the test and control groups (**Figure 3G**). Lastly, the levels of FSH remained elevated throughout the study, increasing from ∼5-10ng/mL(*31*) immediately after ovariectomy and reaching an average of 42ng/mL in both groups in the 20-week cohort (**Figure 3H**). On the 18th week post implantation, a downward trend in FSH was evident in both groups. This downward trend of FSH suggests that the levels of circulating estradiol and inhibins reached the threshold required to restore the negative feedback on the HPG axis.

### 3.4 Antral follicles were recovered in non-encapsulated and encapsulated human ovarian tissue grafts

Histological analysis of retrieved grafts confirmed the presence of follicles at different development stages in the implanted tissue. In all the retrieved samples, multiple primordial and primary follicles were present in each time point for control and test groups (**Supplemental Figure 2**).

Additionally, we identified three antral follicles in the non-encapsulated group, one in the 15-week cohort (800µm in diameter), and two in the 20-week cohort (1.5 mm and 2.5 mm in diameter). One antral follicle was observed in the encapsulated group in the 20-week cohort, measuring 3.5 mm in diameter, validating that follicles can reach antral stages within the capsule and without direct vascularization. The morphology of the antral follicles in the retrieved human xenografts matched the size and composition of a normal antral follicle (**Figure 4A**). Each antral follicle had a clear antral cavity surrounded by an organized layer of mural granulosa cells, separated from a well-defined theca layer by a basement membrane (**Figure 4B-E**). Similar to the non-encapsulated tissue, follicles that developed to an antral stage in the encapsulated tissue had an antral cavity, lined with granulosa cells separated by a basement membrane from the theca layer (**Figure 4F-H).** As expected, the non-encapsulated graft contained visible blood vessels with red blood cells in them, suggesting revascularization of the grafted tissue by the host vasculature. As expected and desired, the encapsulated tissue remained isolated from the host vasculature. Despite the absence of revascularization, critical for prevention of immune rejection, the antral follicles in the encapsulated tissue had a similar organization of granulosa and theca layers (**Figure 4G, H**), suggesting normal folliculogenesis and steroidogenesis.

**Figure 4:**
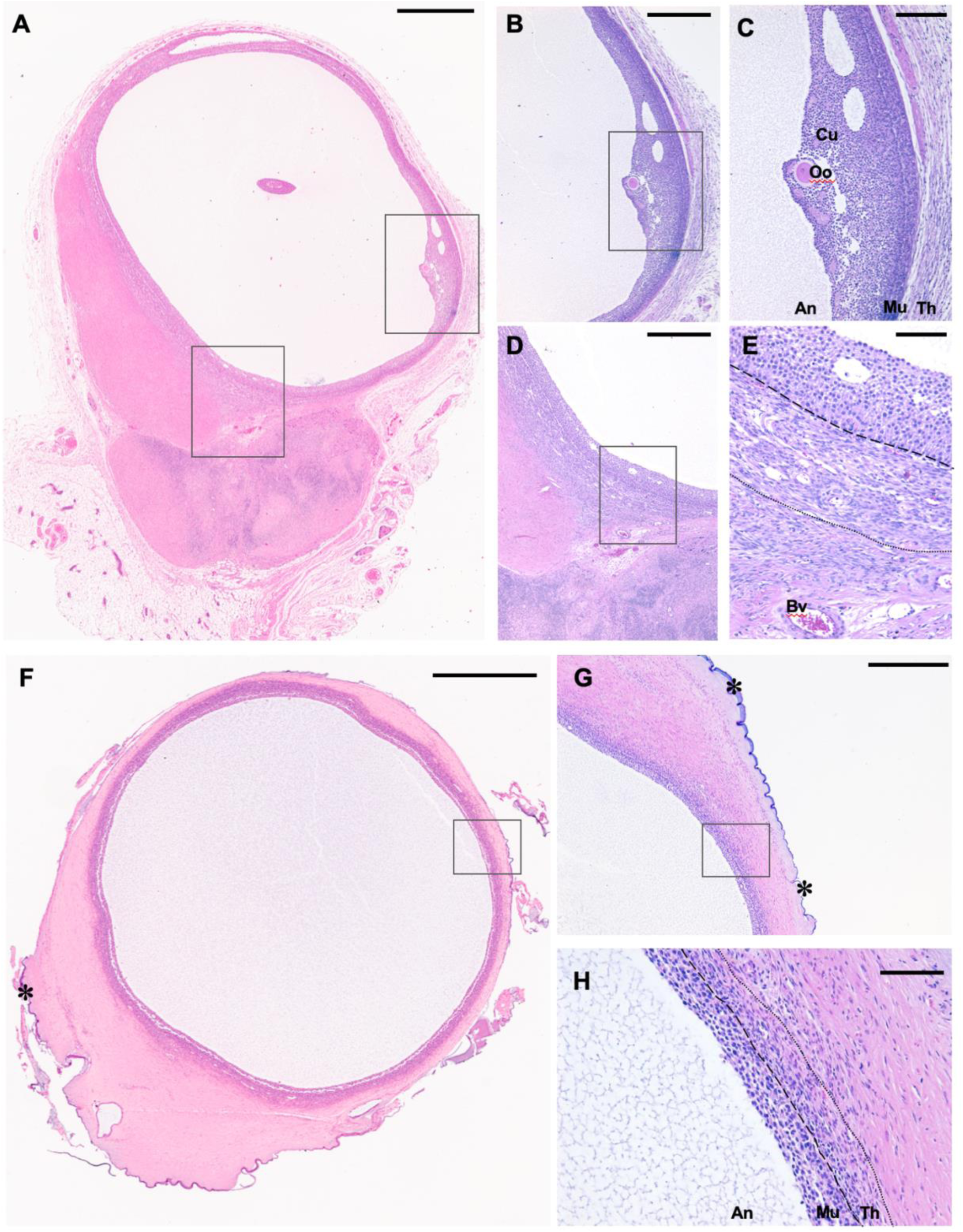
Antral follicles from non-encapsulated (A-E) and encapsulated (F-H) tissue after 20 weeks of implantation. Scale bar represents 1mm (A, F), 500pm (B, D, G), or 100pm (C, E, H). Oo: oocyte, Cu: cumulus granulosa cells, An: antrum, Mu: mural granulosa ceils, Th: theca layer, Bv: blood vessel, PEG capsule.

## Discussion

Utilizing ovarian cortex from 8 young donors, this study provides proof of concept that the hydrogel-based immune-isolating capsule is successful in sustaining folliculogenesis and steroidogenesis comparable to that achieved with non-encapsulated grafts, offering a possible cure for cancer patients with POI. Taken into consideration the inter- and intra-donor heterogeneity in follicle density, we took an extra care in forming the encapsulated and non-encapsulated groups such that donor age, our best predictor for ovarian follicle density and reserve were not significantly different between groups. Despite the expected heterogeneity in follicular density in human donors, all ovariectomized mice implanted with encapsulated or non-encapsulated tissue experienced vaginal opening, at least one estrus cycle indicative of appropriate integration of feedback systems from the ovarian implant. Equally good performance of the encapsulated and non-encapsulated tissue in the number of cycles achieved indicates that the hydrogel-based immune-isolating capsule did not prevent endogenous mouse gonadotropin stimulation of the encapsulated human ovarian cortex and ease of hormone diffusion. This is further emphasized by the similar and not statistically different levels of estradiol achieved in ovariectomized mice with encapsulated and non-encapsulated human ovarian tissues.

The hormonal and cycle dynamic information obtained in this study helps draw a picture of how the human ovarian tissue is integrating within the mouse HPG axis. Intact mice experience estrus every 4 days(*25, 28*), which correlates with the duration of murine follicle development where it takes 4 days for the small antral follicles to reach preovulatory follicle stages and 18 days to complete folliculogenesis from primordial stage to ovulation(*32*). In contrast, human follicles can take up to 180 days to grow from primordial to ovulatory stage and approximately 3 months to reach 1-2 mm in diameter(*29*). In our study, ovariectomized mice were implanted with cryopreserved human ovarian cortical tissue containing only primordial and primary follicles. As expected, the first changes in vaginal opening and estrous cyclicity appeared 6 weeks after implantation, suggesting presence of growing follicles and matching the timeline of human folliculogenesis of about 1.5-2 months needed for human follicles to reach preantral and antral stages (*29*). Interestingly, the cycle length in ovariectomized mice implanted with human ovarian tissue extended to weeks in our study, confirming that despite being stimulated by the endogenous host-produced gonadotropins human follicles followed their internal biological clock. Mice demonstrated variability in cycle duration and frequency (all variations shown in **Supplemental Figure 1**), with some having multiple cycles every few weeks and some mice only experiencing a few cycles. The mouse-to-mouse variation is likely due to the inevitable heterogeneity of follicular pool in the human ovarian tissue graft.

Restoration of ovarian endocrine function was also confirmed by the increased levels of estradiol beginning in the sixth week post-implantation, and overall increase in estradiol levels at a cohort level. There was a trend of increasing estradiol levels positively correlating with the time post-implantation progressed, which was expected as the growing follicles needed time to mature. The additional metric we analyzed was FSH levels, which remained elevated throughout the study, but began to trend downward starting at 18 weeks, suggesting that the levels of estradiol and inhibins reached sufficient levels and remained elevated for long enough to restore the negative feedback on the hypothalamus and pituitary. Importantly, our data suggests that mouse gonadotropins stimulated folliculogenesis and steroidogenesis in human ovarian tissue and human hormones influenced the reproductive organs inducing vaginal opening, estrous cyclicity and integration within the HGP axis. Our data agree with results from Man *et al.*, where serum hormone levels were shown to change with tissue implantation after bilateral ovariectomy(*33, 34*). Furthermore, our data also demonstrates that there were no significant differences in hormone production between the non-vascularized, immune-isolated tissue and non-encapsulated, vascularized control grafts with regards to the onset of graft function and longevity. Interestingly, our results of human ovarian tissue grafted in mice also align with the clinical data reporting that it takes approximately 19 weeks for hormone levels to return to physiological levels in human studies(*22*). Lastly, the presence of theca cells in the antral follicles and increased levels of estradiol further confirm normal steroidogenesis and folliculogenesis. According to the two-cell/two-gonadotropin theory, theca cells that convert cholesterol into androstenedione must be present for granulosa cells to make high levels of estradiol(*35*).

Despite the promising results, our study had a few limitations. First, due to the nature of ovarian follicles and thickness of the human cortex it was impossible to analyze the follicle density without damaging the tissue before implantation. As a result, we were unable to say with certainty how many follicles were implanted in each mouse. A survival dye, such as Neutral red, could be used for follicle counts; however, it would require thinner pieces of tissue which would increase the number of follicles activated due to mechanical stress and excessive manipulation. Follicle activation is of critical concern as too many activated follicles would shorten graft longevity. As a compromise, we minimized the extent of human tissue manipulation, used donor’s age as the determining factor for extrapolating follicle density, and utilized a mixture of cortical tissue such that the average donor age between both groups was similar.

Second, we carried out our study for 20 weeks, attempting to match the duration of the study with human folliculogenesis. Human cortex contains mostly primordial follicles that would only reach antral stages after 20 weeks. As expected, we observed the early signs of endocrine function, including vaginal opening, estrous cyclicity, measurable systemic levels of estradiol, and a downward trend in FSH levels. This duration is in line with the data reported by Khattak et al(*22*) in human studies, but a longer study would allow us to reach complete restoration of the HPG axis function, possibly several waves of activation, and even determine how long it would take to exhaust the implanted follicle reserve and establish a dose-response relationships. Third, while we have been able to measure estradiol and FSH in blood serum, we were not able to collect enough blood to test for multiple hormones, such as progesterone, AMH and inhibins, due to the limited blood sampling in a mouse model, and we therefore only report estradiol levels at terminal time points. The ability to collect more blood serum or use test kits that are more sensitive and require less blood would allow us to have a more nuanced picture of what was happening with multiple hormones over the duration of this study and their cyclic profiles.

Lastly, we used immunocompromised mice in this study which did not allow us to test the immune-isolating ability of the capsules. Yet, this study still provides important evidence that despite encapsulation and lack of direct revascularization, human ovarian tissue grafts have survived and functioned, relying solely on diffusion. The next step would be to evaluate the capsules in an immune competent mouse model that would inform us on capsule’s ability to withstand immune rejection and follicle expansion without fail.

In conclusion, we have shown that both encapsulated and non-encapsuled large human antral follicles grow in an ovariectomized NSG mouse model. The grafts produced estradiol, decreased FSH and allowed estrous cyclicity. These results indicate that the graft interfaced with the mouse hypothalamus/pituitary axis, but also that encapsulation did not preclude hormone diffusion or follicle development.

## 4. Author Contributions

Conceptualization and funding: AS

Study design: MAB, VP, AS

Murine experiments: MAB, MAW, DS

Histological sample preparation: MAB, ML, AT, BR

Follicle counting and figure compilation: JHM, GB, MB

Wrote original manuscript draft: MAB, AS

Manuscript review & editing: MAB, MC, VP, AS

## Funding

National Institutes of Health grant R01HD104173, R01EB022033 (MAB, MC, AS)

National Institutes of Health grant T32DE007057 (MAB)

National Institutes of Health grant T32 HD079342 (MAB)

National Institutes of Health Z1A HD009005; Eunice Kennedy Shriver NICHD (MAB)

Musculoskeletal Transplantation Foundation (MTF) Biologics Research Grant Application 2021

Eunice Kennedy Shriver NICHD/NIH grant R24HD102061(UVA Ligand Assay and Analysis Core)

## Acknowledgments

Figures were created with BioRender. Hormone analysis was conducted by the University of Virginia Center for Research in Reproduction Ligand Assay and Analysis Core. Samples were processed for sectioning by the University of Michigan Dental Histology Core. Whole slide scanning provided by the University of Michigan ULAM Pathology Core.

---

**Supplemental Figure 2:**
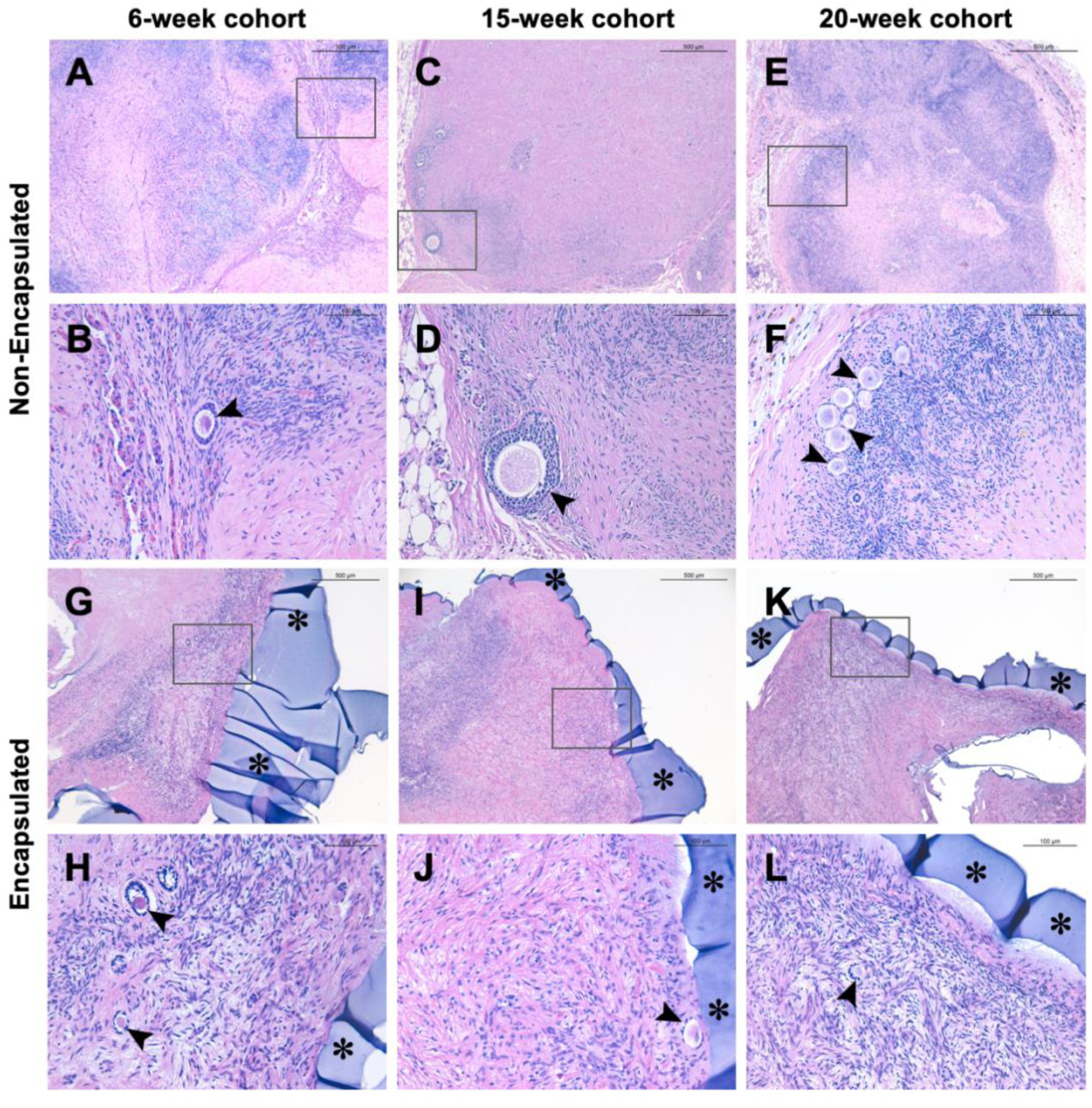
Representative histological images from non-encapsulated (A-F) and encapsulated (G-L) tissues. Scale bar represents 500um (A,C,E,G,I,K) or 100um (B, D, F, H, J, L). *: PEG capsule, ►: follicle.

**Supplemental Table 1:** Donor Information.

| Donor ID | A | B | C | D | E | F | G | H | Average ± Std. Dev. |
| --- | --- | --- | --- | --- | --- | --- | --- | --- | --- |
| Age (years) | 18 | 16 | 25 | 27 | 18 | 25 | 31 | 28 | 23.5 ± 5.5 |
| Height (cm) | 165 | 155 | 173 | 173 | 163 | 180 | 152 | 157 | 164.8 ± 9.9 |
| Weight (kg) | 48.8 | 51.1 | 61.7 | 86.5 | 89 | 95 | 87.2 | 56 | 71.9 ± 19.3 |
| Body Mass Index (kg/m<sup>2</sup>) | 17.9 | 21.3 | 20.6 | 28.9 | 33.5 | 29.2 | 37.7 | 22.7 | 26.5 ± 6.9 |
| Right Ovary Volume (mL) | 6 | 2.5 | 9 | 5 | 9 | 12 | 11 | 5 | 7.4 ± 3.3 |
| Left Ovary Volume (mL) | 5 | 5 | 12 | 6 | 7 | 10 | 9 | 10 | 8.0 ± 2.6 |
| Cold Ischemic Time* (hours) | 23.7 | 15.4 | 13.7 | 21.2 | 14.7 | 25.5 | 27.7 | 20.4 | 20.3 ± 5.3 |
| Cardiac arrest/downtime (minutes) | YES | YES<br>20 | YES<br>50 | NO | NO | YES<br>15 | YES<br>0 | YES<br>19 |  |
| Cross-clamp Date and Time** | 5/21/21<br>10:42 | 5/25/21<br>6:58 | 5/29/21<br>10:19 | 7/20/21<br>18:54 | 7/27/21<br>8:50 | 1/2/22<br>13:49 | 1/14/22<br>17:26 | 1/14/22<br>17:26 |  |
| Ethnicity | W | W | W | B | W | W | W | W |  |
| Cause of Death | Anoxia | Anoxia | Anoxia | CV/S | Anoxia | Anoxia | Anoxia | Anoxia |  |
| Storage Solution | NS | NS | NS | UW | NS | NS | UW | NS |  |

B, Black or African American; W, White.
CV/S, cardiovascular/stroke
UW, University of Wisconsin Solution; NS, Normal Saline.
\*Time at which the organ is cut from blood/oxygen supply.
\*\*Time between cross-clamp and the beginning of tissue processing

**Supplemental Table 2:** Donor information by mouse.

| Group | Time Point | Mouse | Donors |
| --- | --- | --- | --- |
| Encapsulated | 6 weeks | 3 | C, F |
|  |  | 10 | B, E, F |
|  |  | 20 | E, G |
|  |  | 28 | E, G |
|  | 15 weeks | 11 | B, F |
|  |  | 15 | A, B, F |
|  |  | 21 | G, H |
|  |  | 27 | G |
|  | 20 weeks | 5 | A, F |
|  |  | 8 | F, G |
|  |  | 12 | E, F |
|  |  | 13 | F |
|  |  | 17 | G |
|  |  | 22 | G |
|  |  | 25 | E, G |
| Non-Encapsulated | 6 weeks | 1 | A, F |
|  |  | 14 | E, F |
|  |  | 18 | E, G, H |
|  |  | 24 | E, G, H |
|  | 15 weeks | 4 | C, F |
|  |  | 6 | B, F |
|  |  | 19 | G, H |
|  |  | 23 | E, G, H |
|  | 20 weeks | 2 | B, C, G |
|  |  | 7 | F, G |
|  |  | 9 | B, E, F |
|  |  | 16 | A, F |

